# Time-resolved single-cell sequencing identifies multiple waves of mRNA decay during mitotic exit

**DOI:** 10.1101/2021.04.17.440266

**Authors:** Lenno Krenning, Stijn Sonneveld, Marvin E. Tanenbaum

**Affiliations:** Oncode Institute, Hubrecht Institute – KNAW and University Medical Center Utrecht, Utrecht, the Netherlands; Division of Cell Biology, Oncode Institute, Netherlands Cancer Institute, Amsterdam, the Netherlands

## Abstract

Accurate control of the cell cycle is critical for development and tissue homeostasis and requires precisely-timed expression of many genes. Cell cycle gene expression is regulated through transcriptional and translational control, as well as through regulated protein degradation. Here, we show that widespread and temporally-controlled mRNA decay acts as an additional mechanism for gene expression regulation during the cell cycle. We find that two waves of mRNA decay occur sequentially during the mitosis-to-G1 phase transition, and identify the deadenylase CNOT1 as a factor that contributes to mRNA decay during this cell cycle transition. Collectively, our data show that, akin to protein degradation, scheduled mRNA decay helps to reshape cell cycle gene expression as cells move from mitosis into G1 phase.

## Introduction

Cell division is essential for the development and homeostasis of multicellular organisms. Precise control over cell division is paramount, as errors may contribute to carcinogenesis (Hanahan and Weinberg, 2011; Malumbres and Barbacid, 2001). In order to divide, cells pass through a number of different phases collectively referred to as the cell cycle. The cell cycle in somatic cells consists of four phases: (1) in G1 phase a cell grows and prepares for DNA replication; (2) in S phase the DNA is replicated; (3) in G2 phase a cell prepares for segregation of the replicated genome; (4) in M phase (or mitosis) the cell divides and then enters into G1 phase of the next cell cycle. Progression through the cell cycle is accompanied by the periodic expression of many genes (referred to as cell cycle genes), whose protein products are required in a particular cell cycle phase (Bar-Joseph et al., 2008; Chaudhry et al., 2002; Cho et al., 1998, 2001; Grant et al., 2013; Whitfield et al., 2002). Deregulated expression of cell cycle genes can decrease the fidelity of cell division. For instance, reduced expression of G2 and M phase cell cycle genes impedes mitotic entry and affects the fidelity of chromosome segregation (Laoukili et al., 2005). Conversely, a failure to suppress expression of G2 and M phase genes as cells enter G1 phase results in a shortened G1 phase and causes DNA replication errors (García-Higuera et al., 2008; Park et al., 2008; Sigl et al., 2009), and can even contribute to carcinogenesis (Bortner and Rosenberg, 1997; Coelho et al., 2015; Kalin et al., 2006; Kim et al., 2006; Vaidyanathan et al., 2016). These examples highlight the importance of tightly controlled gene expression for proper execution of the cell cycle.

To restrict cell cycle gene expression to the correct cell cycle phase, cells need to activate, but also to repress the expression of cell cycle genes as they move from one phase to the next. Scheduled protein degradation plays an important role in repression of cell cycle gene expression by ensuring that protein expression is restricted to the appropriate cell cycle phase (Nakayama and Nakayama, 2006; Vodermaier, 2004). In addition, cells prevent *de novo* synthesis of proteins through inhibition of transcription to further restrict protein expression to the correct cell cycle phase (Bertoli et al., 2013; Sadasivam and DeCaprio, 2013). While inhibition of transcription will eventually lower mRNA levels and thus decrease protein synthesis rates, this process is relatively slow, as it requires turnover of the existing pool of mRNAs. To circumvent this, cells can shut down translation or degrade pre-existing transcripts when transitioning from one cell cycle phase to another. Indeed, control of mRNA translation also contributes to the regulation of gene expression during the cell cycle (Kronja and Orr-Weaver, 2011) and several hundreds of genes are subject to translational regulation at different phases of the cell cycle (Stumpf et al., 2013; Tanenbaum et al., 2015).

Regulation of mRNA stability during the cell cycle has been studied relatively little, but recent work suggests that this type of regulation also contributes to restriction of cell cycle gene expression. Dynamic changes in mRNA stability during the cell cycle were observed in yeast using fluorescent in situ hybridization (FISH). More specifically, cyclin B mRNA was shown to be destabilized upon completion of mitosis (Trcek et al., 2011). Globally, mRNA synthesis and decay rates during the cell cycle of yeast were derived through metabolic mRNA labeling in synchronized populations, resulting in the identification of several hundred genes that show periodic changes in mRNA synthesis and degradation rates (Eser et al., 2014). Regulation of mRNA stability is also reported to occur during the human cell cycle. For instance, the transcription factor ERG was shown to control the degradation of a set of mRNAs during S phase (Rambout et al., 2016). Recently, global mRNA synthesis and degradation rates during the human cell cycle were determined (Battich et al., 2020). Here, a newly-developed method that simultaneously quantifies metabolically labeled and preexisting unlabeled transcripts in individual cells was used to determine synthesis and degradation rates of individual transcripts along the cell cycle. Together, these studies demonstrate that the stability of many mRNAs change during the cell cycle. However, due to the relatively long measurement time required for the pulse-chase approach used by Battich et al. (up to 6 hours), accurate dynamics and rapid changes, especially around the transition points in the cell cycle, are difficult to determine.

To obtain a highly quantitative view of transcriptome dynamics during cell cycle phase transitions, we set up a method that combines singe cell mRNA sequencing and live-cell imaging of cell cycle progression to map transcriptome-wide mRNA expression levels with high temporal resolution during the cell cycle. We focus specifically on the mitosis to G1 (M-G1) phase transition when cells divide and enter into a new cell cycle, as gene expression needs to be ‘reset’ after cell division. The widespread protein degradation that occurs during the M-G1 phase transition is thought to contribute to this reset (Castro et al., 2005; Harper, 2002; Peters, 2002; Vodermaier, 2004). We hypothesized that, analogous to scheduled protein degradation, mRNA decay might play an important role in resetting cell cycle gene expression by limiting the carry-over of pre-existing G2/M-specific transcripts from one cell cycle into the next. Using our method, we identified two temporally-distinct waves of mRNA decay: the first wave is initiated during mitotic exit and the second wave is initiated within the first hours of G1 phase. For several of these genes, we show that mRNA decay requires CNOT1, a subunit of the CCR4-NOT mRNA deadenylase complex that shortens the poly(A) tail of mRNAs, generally resulting in their decay (Garneau et al., 2007; Yamashita et al., 2005). Together, our findings demonstrate that, analogous to protein degradation, mRNA degradation occurs at the M-G1 phase transition, and provides an important contribution to the reset of the transcriptome after cell division.

## Results

### Time-resolved transcriptome profiling during the cell cycle using the FUCCI system

To obtain a detailed view of mRNA levels as cells progress from M phase into G1 phase, we developed a method that connects live-cell microscopy with single cell RNA sequencing (scRNA-seq), through fluorescence activated cell sorting (FACS). This method allows us to assign an accurate, ‘absolute’ cell cycle time (i.e., the time in minutes since G1 phase entry) to individual sequenced cells, which we used to generate a high-resolution, time-resolved transcriptome profile of the M-G1 phase transition.

To assign an absolute cell cycle time for each cell, we expressed the fluorescent, ubiquitination-based cell cycle indicator (FUCCI) system in a human untransformed cell line, RPE-1 (RPE-FUCCI). In the FUCCI system an orange fluorescent protein (FUCCI-G1) is expressed in G1 and early S phase cells, while a green fluorescent protein (FUCCI-G2) is expressed in late S, G2 and early M phase (Fig. 1A) (Sakaue-Sawano et al., 2008). Importantly, the expression levels of both fluorescent markers change over time within each cell cycle phase, potentially allowing precise pinpointing of the cell cycle time of individual cells based on the fluorescence intensity of the FUCCI reporter. We used live-cell microscopy to measure FUCCI-G1 fluorescence intensity in cells as they progressed through G1 phase and observed a monotonic increase during the first 6–8 hours after G1 entry (Fig. S1A). To allow accurate calculation of a cell cycle time based on FUCCI-G1 fluorescence intensity, we fit the average FUCCI-G1 fluorescence intensity to a polynomial equation (Fig. 1B and Supplementary table 1). Using this equation, a cell cycle time can be calculated for each fluorescence intensity of the FUCCI-G1 marker as assayed by time-lapse microscopy. However, in the scRNA-seq protocol the fluorescence intensity of each sequenced cell is measured by FACS analysis. To compare fluorescence intensities measured by FACS and imaging, we normalized FUCCI-G1 fluorescence. Since early S phase cells can be identified in both live-cell imaging experiments and by FACS analysis (Fig. 1C and S1B), the mean fluorescence intensity of the FUCCI-G1 marker in early S phase can be used as a normalization factor to directly compare the FUCCI-G1 fluorescence intensity values obtained by imaging and FACS (Fig. 1C, S1B; see Methods). Using this normalization factor and the fluorescence intensity of the FUCCI-G1 marker as assayed by FACS, it is possible to map individual G1 cells assayed by FACS onto time-lapse microscopy data, allowing us to pinpoint the precise cell cycle time of each cell that is sorted by FACS.

**Figure 1.**
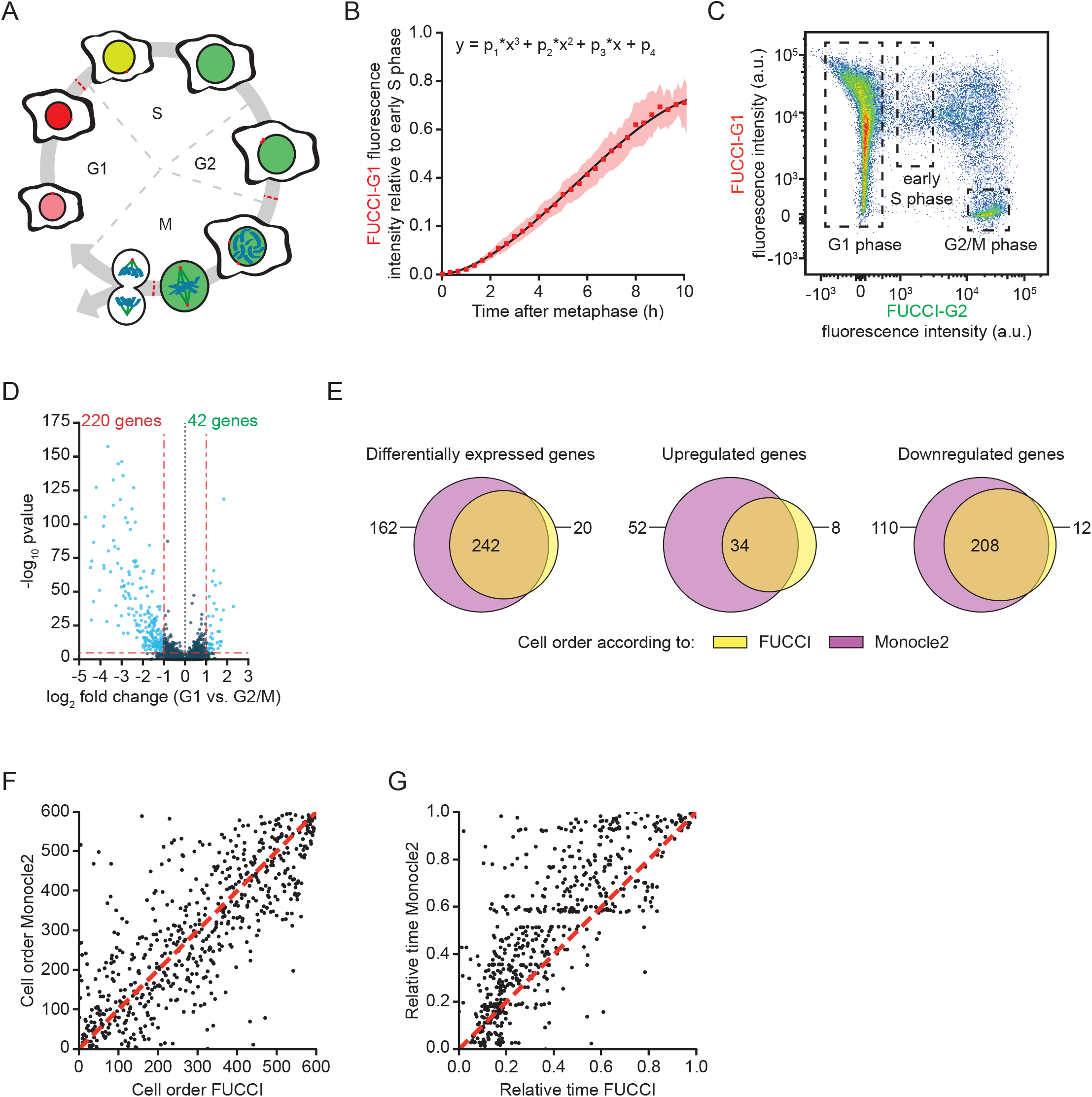
A method for time-resolved transcriptome analysis during the cell cycle. **A)** Schematic representation of the FUCCI-system. **B)** Modeling of FUCCI fluorescence intensities. Asynchronously growing RPE-FUCCI cells were analyzed by live-cell imaging (Fig. S1A). Subsequently, FUCCI-G1 fluorescence intensities were measured and normalized to the average fluorescence in early S phase cells (see methods). Squares and shading represent the mean fluorescence and SEM of three individual experiments. The mean FUCCI-G1 marker fluorescence was fit to a third-order polynomial (black line, equation above plot). The fit has no biological meaning, but serves to approximate the data to allow calculation of estimated fluorescence intensities at different time points during the cell cycle. **C)** FACS analysis of asynchronously growing RPE-FUCCI cells, including the gating strategy used for the identification and isolation of G1, early S and G2/M phase cells. **D)** Differential gene expression analysis of RPE-FUCCI cells in G2/M-versus G1 phase. **E)** Venn diagram comparing differentially expressed genes (both up- and downregulated in G1 versus G2/M phase) identified after FUCCI- or Monocle2-based cell ordering. **F)** Comparison of FUCCI and Monocle2 based ordering of G1 phase cells. Dashed line indicates identical order of cells. **G)** Comparison of G1 phase timing based on either FUCCI-based ordering or pseudo-timing based on trajectory inference by Monocle2. Dashed line indicates identical timing of FUCCI and Monocle2.

To validate that our method of converting FACS fluorescence intensities into absolute cell cycle times is accurate, we performed a control experiment. We blocked cells in mitosis using the microtubule stabilizing drug Taxol for various durations, preventing entry of cells in G1 phase. For cells already in G1 phase the FUCCI-G1 fluorescent signal continues to increase. As no new cells enter G1 phase, a gradual loss of cells with low FUCCI-G1 fluorescence is observed by FACS (Fig. S1C). By mapping the population of cells that is lost after different times of Taxol treatment we could calculate the FUCCI-G1 fluorescence intensity associated with cells that had spent various times in G1 phase. Comparison of this method to the values obtained with the polynomial equation revealed very similar results (Fig. S1D). Thus, we conclude that we can accurately determine the time a cell has spent in G1 phase based on its FUCCI-G1 fluorescence as measured by FACS.

To identify changes to the transcriptome throughout the M-G1 phase transition, we FACS-isolated single G2, M and G1 phase cells based on their FUCCI-G1 and FUCCI-G2 fluorescence (Fig. 1C), and subjected them to scRNA-seq. In total, 1152 cells were sequenced in three replicate experiments, of which 841 cells passed quality checks (see Methods) and were used to generate a high-resolution temporal transcriptome profile of the M-G1 phase transition. Since the FUCCI system does not discriminate between cells in G2 and M phase, and as there are few transcriptome changes between these two phases (Tanenbaum et al., 2015), we averaged the transcript levels of all cells in G2 and M phase (referred to as G2/M). The average G2/M expression levels of individual genes displayed a high correlation between different replicate experiments (Fig. S1E-G), allowing us to pool the data from the different experiments. The final dataset consisted of 86 G2/M phase cells and 755 cells from various time points in G1 phase (up to 9 hr after the M-G1 phase transition) (Fig. S1H and Supplementary table 1). After initial data processing (see Methods), we performed differential transcriptome analysis comparing G2/M phase to G1 phase cells. This analysis identified 220 genes that were downregulated and 42 genes that were upregulated when cells progressed from G2/M phase into G1 phase (using a cutoff of >2-fold expression change and a p-value of <10^−5^) (Fig. 1D and Supplementary table 1). Gene ontology analysis revealed that these differentially expressed genes were strongly enriched for cell cycle functions, as expected (Fig. S1I).

To compare our method of cell cycle time determination with previous computational methods, we used Monocle2, an *in silico* trajectory inference method that orders cells based on their transcriptomes (Qiu et al., 2017b, 2017a; Trapnell et al., 2014). We aligned cells using trajectory inference (Fig. S1J and see Methods), and subsequently performed differential transcriptome analysis, which identified 318 downregulated genes and 86 upregulated genes in G1 phase compared to G2/M phase (Fig. S1K and Supplementary table 1). There was a large overlap between the differentially expressed genes identified by Monocle2 and our FUCCI-based method (Fig. 1E), and we found a good overall correlation between FUCCI-based ordering and Monocle2-based ordering of G1 phase cells (Fig. 1F and S1J). Monocle2 cannot assign absolute cell cycle times, instead it can compute a ‘pseudo time’ for each G1 phase cell assuming that transcriptome changes occur smoothly over time. Comparing the pseudo time assigned by Monocle2 with the cell cycle time assigned by our FUCCI-based method revealed differences between both methods. In general, Monocle2 computed larger time intervals between cells early in G1 phase compared to our FUCCI-based method (Fig. 1G). As Monocle2 computes the time intervals between cells based on the magnitude of transcriptome changes, a possible explanation for this observation is that transcriptome changes are larger in early G1 phase than at the end of G1 phase, and Monocle2 thus positions cells in early G1 phase too far apart in (pseudo) time. In conclusion, by using the FUCCI-based single-cell sequencing approach we could generate a high-resolution, time-resolved transcriptome profile of cells spanning the transition from M phase into G1 phase.

### mRNA levels decline in multiple waves after cell division

As discussed above, we found a large group of genes (220) for which mRNA levels decline at the M-G1 phase transition. To determine the precise moment when mRNA levels started to decline for each gene, we fit the data for individual mRNAs to a smoothing spline and determined the moment of maximum negative slope of the spline, which is the moment when the mRNA level declined most rapidly (referred to as spline analysis; see Methods). Strikingly, the decline in mRNA levels of various genes initiated at two distinct times in the cell cycle: the first occurred around the time of mitotic exit and the second at ∼80 minutes into G1 phase (Fig. 2A and Supplementary table 1). To examine these two ‘waves’ of mRNA decline in more detail, we divided the 220 mRNAs into two groups: one for which the maximum negative slope occurred during mitotic exit (*immediate decrease*) and one for which the maximum negative slope occurred during G1 phase (*delayed decrease*) (Supplementary table 1, see Methods). Plotting the average slope over time for both groups (Fig. 2B) revealed that the mRNAs in the *immediate decrease* group declined most rapidly during the M-G1 phase transition and continued to decline during the first 2–3 hours of G1 phase, whereas the mRNAs in the *delayed decrease* group mostly declined between 1–4 hours after the start of G1 phase. For both groups, the slopes of individual mRNAs were mostly centered around zero at later times (>8 hours) in G1 phase, suggesting that most mRNAs reached a new steady-state level at later time-points in G1 phase.

**Figure 2.**
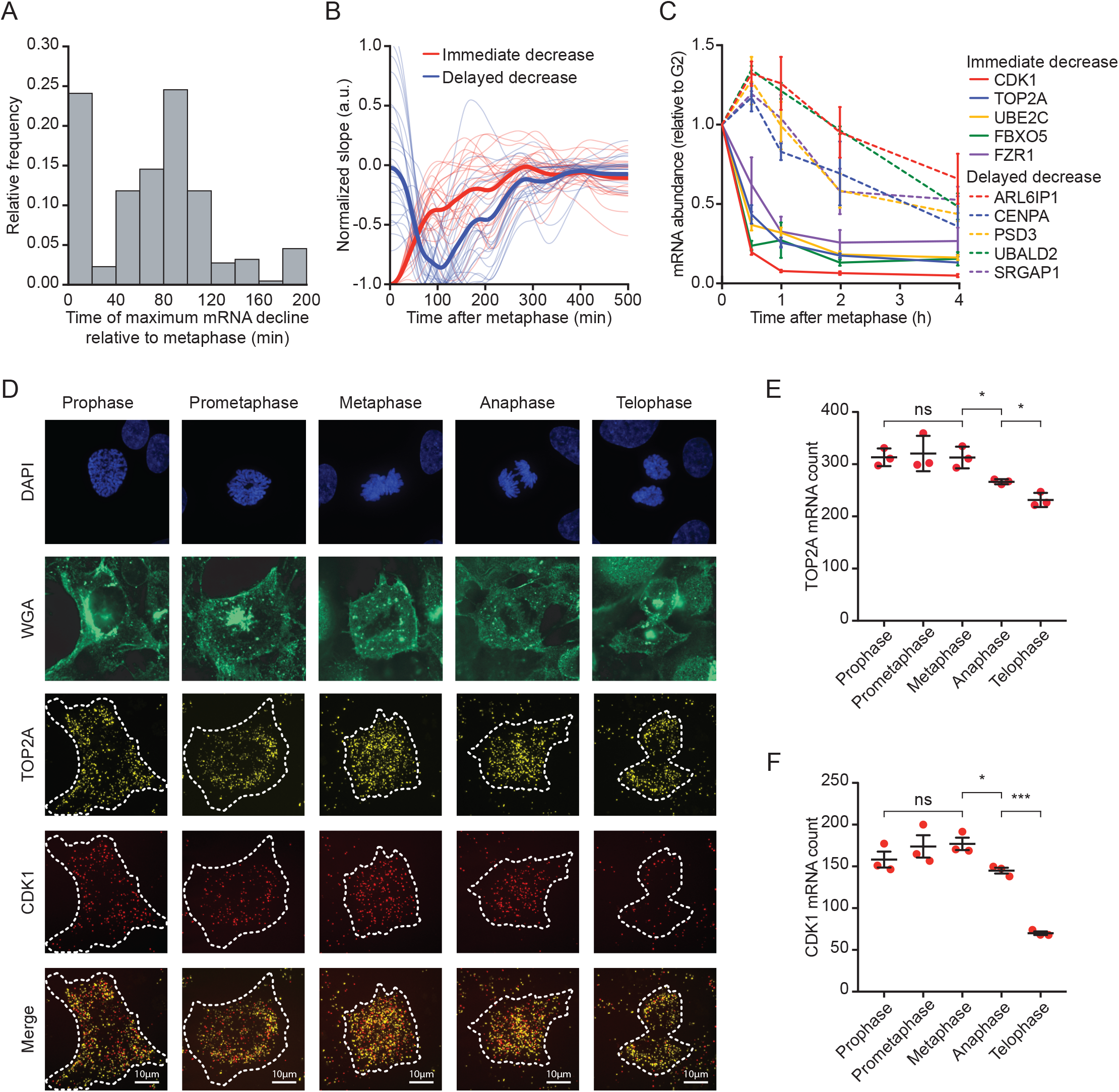
Reduction in mRNA levels occurs in multiple waves, during and after cell division. **A)** Time, relative to metaphase, of the highest rate of mRNA decrease for 220 downregulated genes (see Methods). **B)** Average slope of mRNA levels over time for genes that display immediate (thick red line) or delayed decrease (thick blue line). Thin red and blue lines show a random selection of 25 individual genes belonging to the *immediate* or *delayed decrease* group, respectively. **C)** Validation of different waves of mRNA reduction. RPE-FUCCI cells at different stages of the cell cycle were isolated by FACS based on FUCCI fluorescence (see Fig. S2 for gating strategy). mRNA expression levels of indicated genes was measured by RT-qPCR. Five genes from the *immediate decrease* group and five genes from the *delayed decrease* group were selected. Note that the moment of decrease as measured by RT-qPCR closely mirrors the values obtained by modeling of our single cell sequencing data (see Supplementary table 1). Lines with error bars represent average ± SEM of 3 experiments. **D)** Example images of TOP2A and CDK1 mRNA levels during the different stages of mitosis. Asynchronously growing RPE-1 cells were fixed and stained for DNA (DAPI), membranes (WGA) and TOP2A and CDK1 mRNA (using bDNA-FISH). Scale bar, 10 µm. **E-F)** Quantification of TOP2A (E) and CDK1 (F) transcripts (shown in panel D) using ImageJ. Each dot represents a single experiment and lines with error bars represent average ± SEM of 3 experiments (at least 14 cells per experiment per mitotic phase, see Supplementary table 2). P-values are based on a one-tailed unpaired Student’s t-test, and are indicated as * (p<0.05), ** (p<0.01), *** (p<0.001), ns = not significant.

To confirm that mRNA levels decline in two distinct temporal waves, we used RT-qPCR to measure mRNA levels for five genes in the *immediate decrease* group (CDK1, TOP2A, UBE2C, FBXO5, and FZR1) and five genes in the *delayed decrease* group (ARL6IP1, CENPA, PSD3, UBALD2, and SRGAP1) in G2/M phase and at various time-points in G1 phase (Fig. S2). Consistent with the RNA sequencing data, we observed two distinct waves of mRNA decline by RT-qPCR (Fig. 2C). The minor increase in mRNA levels seen for the 1 hr time-point in the *delayed decrease* group is likely an artifact caused by comparing a highly synchronized population of early G1 phase cells (which have not yet initiated the decline of delayed genes and thus express the highest possible levels of these transcripts) to a somewhat more heterogeneous population of G2/M phase cells. Collectively, these data demonstrate that there are two distinct waves of mRNA decline during the M-G1 phase transition.

To determine the moment of mRNA decline more precisely for the *immediate decrease* group, we assessed mRNA levels by single molecule FISH (smFISH) and fluorescence microscopy during different mitotic stages. We fixed asynchronous cultures of cells and stained them specifically for two mRNAs from the *immediate decrease* group (TOP2A and CDK1). We focused on TOP2A and CDK1 as they showed a strong mRNA decline after metaphase of mitosis (Fig. 2C). To determine the mitotic stages and the outline of the individual cells, we stained the DNA with DAPI, and the membranes with fluorescent wheat germ agglutinin (Fig. 2D). Quantification of TOP2A and CDK1 mRNA levels at various stages of mitosis revealed a significant decrease in mRNA levels as early as anaphase, and a further decrease in telophase for both genes (Fig. 2E-F). We conclude that the first wave of mRNA decline initiates at the start of anaphase whereas the second wave initiates during early G1 phase.

### mRNA decay drives transcriptomic changes during the M-G1 phase transition

The decline in mRNA levels during early G1 phase may be caused by changes in the rate of mRNA synthesis (transcription) and/or degradation (mRNA stability). To investigate whether mRNA stability is altered during the M-G1 phase transition, we calculated the degradation rate of individual mRNAs using a simple mathematical model (Fig. 3A, see methods). Briefly, our model describes two phases in the mRNA levels over time: in the first phase mRNA levels remain constant (at an initial level of m_0_), in the second phase mRNA levels decline to a new steady state level. The onset of decline is described by t_onset_. The rate of decline is dependent on the mRNA degradation rate (γ), while the new steady-state mRNA level is dependent on the mRNA synthesis rate (µ) and on the mRNA degradation rate (γ). Using a quality of fit analysis (see Methods), we identified the parameters (m_0_, t_onset_, µ, and γ) that resulted in the optimal fit with the data for each of the 220 downregulated genes. Visual inspection showed that the fits described the data well (Fig. S3A-F and Supplementary table 1). Using this approach, we confirmed that the onset of decay for different genes occurred most strongly at two distinct times during the M-G1 phase transition; either during the M-G1 phase transition or during early G1 phase (Fig. S3G), confirming the results from the spline analysis (Fig. 2A).

**Figure 3.**
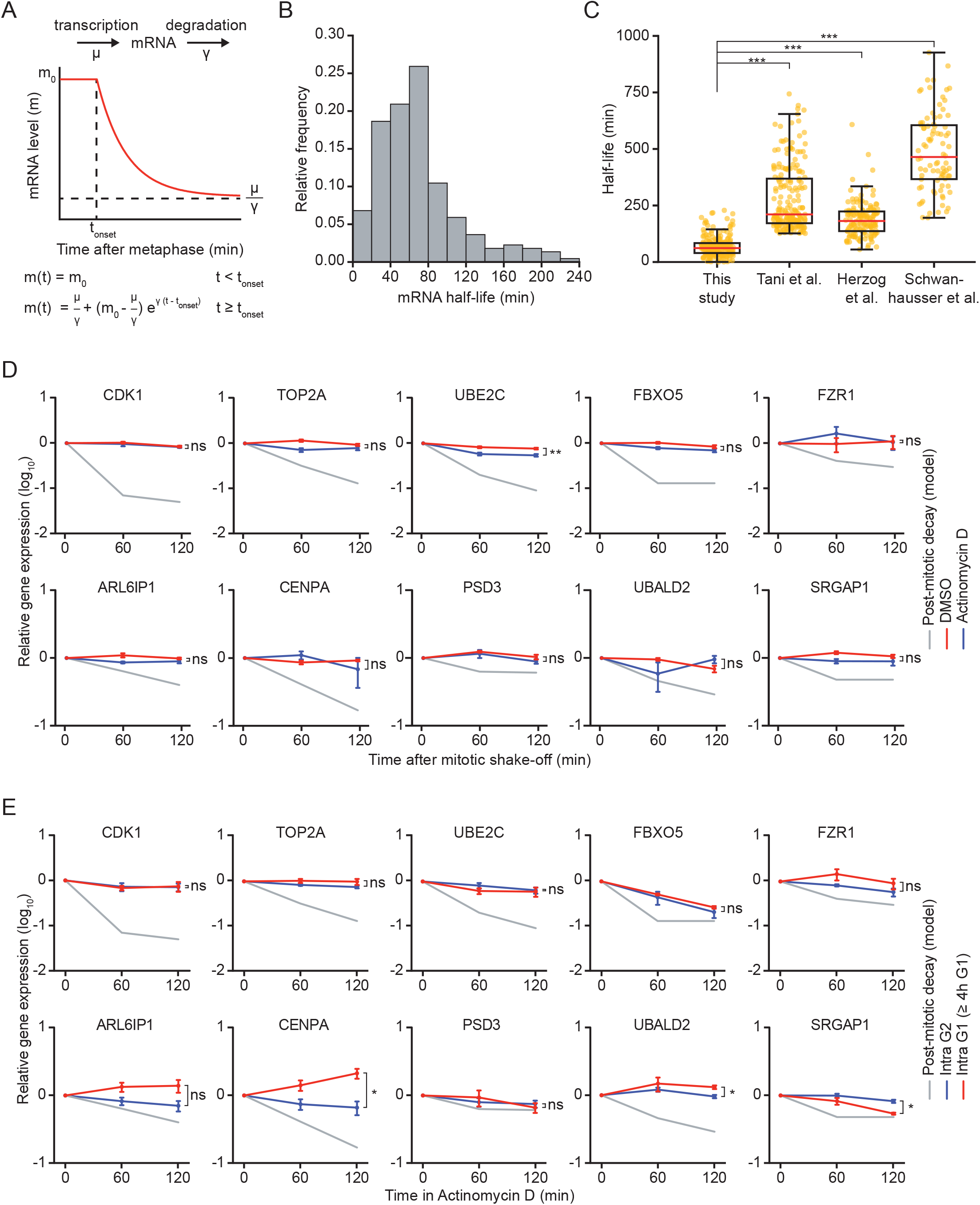
mRNA decay occurs during a brief window of time as cells exit mitosis and enter G1 phase. **A)** Schematic of the mathematical model that was used to fit the decrease in mRNA levels as cells progress from M into G1 phase. **B)** Histogram of mRNA half-lives of the 220 genes that are downregulated during the M-G1 phase transition. **C)** Boxplot of mRNA half-lives of the 220 genes that were found to be downregulated in our data set (This study). mRNA half-lives of the same genes that were measured in previous studies using bulk assays and asynchronous cell populations are also shown; Tani et al. (HeLa cells), Herzog et al. (mouse embryonic stem cells), and Schwanhausser et al. (mouse fibroblasts). **D)** Relative mRNA levels in mitosis after different times of transcription inhibition, as measured by RT-qPCR. Mitotic cells were collected by mitotic shake-off, and cultured for an additional 2 hours in the presence or absence of the transcription inhibitor Actinomycin D (blue and red lines, respectively). For comparison, mRNA levels during the M-G1 phase transition are shown (grey line). Note that mRNA of indicated genes is stable in mitosis, indicating that mRNA is degraded specifically during the M-G1 phase transition. Lines with error bars indicate average ± SEM of 3 experiments. **E)** Relative mRNA levels in G2 and late G1 phase after different times of transcription inhibition, as measured by RT-qPCR. Asynchronously growing RPE-FUCCI cells were treated with Actinomycin D for indicated times. Cells were then FACS-sorted and G2 phase cells and late G1 phase cells (> 4 hr into G1 phase) were isolated based on FUCCI reporter fluorescence. The mRNA level of indicated genes was then measured by RT-qPCR. As in Fig. 3D, mRNA levels during the M-G1 phase transition are shown for comparison (gray lines). Note that mRNA levels are substantially less stable in cells during the M-G1 phase transition compared to G2 or late G1 phase cells. Lines with error bars indicate average ± SEM of 3 experiments. P-values are based on a one-tailed unpaired or paired Student’s t-test (Fig. 3D and 3E or 3C, respectively), and are indicated as * (p<0.05), ** (p<0.01), *** (p<0.001), ns = not significant.

We used mRNA degradation rates extracted from the model to compute the half-lives of the 220 mRNAs that we found to be downregulated in G1 phase. This revealed a median half-life of 61.5 minutes once mRNA levels start to decline during the M-G1 phase transition (Fig. 3B and Supplementary table 1). In most cases the half-lives we computed are substantially shorter than the half-lives of the same mRNAs in asynchronously growing cells as reported previously (Fig. 3C and Fig. S3H-J) (Herzog et al., 2017; Schwanhäusser et al., 2011; Tani et al., 2012). The comparatively short mRNA half-lives we find during the M-G1 phase transition indicate that these transcripts are subject to scheduled degradation. We observed no significant differences between the half-lives of mRNAs belonging to the *immediate decrease* group versus the *delayed decrease* group (Fig. S3K), suggesting that for both groups, mRNA degradation plays an important role in the decline of mRNA levels.

To confirm that mRNAs are subjected to scheduled degradation specifically during the M-G1 phase transition, we examined their stability during mitosis, G2 phase and late G1 phase using an alternative method. To measure mRNA stability in mitosis, we synchronized and arrested RPE-1 cells in prometaphase of mitosis (see Methods), followed by inhibition of transcription for 1 or 2 hours using Actinomycin D. Actinomycin D completely blocked de novo transcription (Fig. S3L) and did not influence the arrest of cells in mitosis (Fig. S3M). None of the ten mRNAs tested (belonging to both the *immediate* and *delayed decrease* groups) showed an appreciable decrease in mRNA levels during the two-hour time window of Actinomycin D treatment, indicating that these mRNAs are much more stable in mitosis than they are during the M-G1 phase transition (Fig. 3D, compare red or blue lines to gray line).

To measure mRNA stabilities in G2 and late G1 phases, we inhibited transcription with Actinomycin D for 1 or 2 hours in asynchronously growing RPE-FUCCI cells. Subsequently, we FACS-sorted populations of G2 cells and late G1 cells (cells that had spent at least 4 hours in G1 phase) and determined mRNA levels of immediate and delayed decay genes with or without Actinomycin D treatment. For all genes tested, mRNA stabilities in either G2 or late G1 phase substantially exceeded the mRNA stability calculated during the M-G1 phase transition (Fig. 3E). Collectively, these data demonstrate that for all genes tested, mRNAs are substantially more stable during G2 phase, mitosis (pre-anaphase), and late G1 phase compared to during the M-G1 phase transition and early G1 phase. Thus, these results uncover an active mRNA decay mechanism that specifically takes place during mitotic exit and early G1 phase.

### CNOT1 stimulates mRNA decay during the M-G1 phase transition

Cytoplasmic mRNA degradation is often initiated by shortening of the poly(A) tail (Eisen et al., 2020), followed by degradation from either end of the mRNA (Garneau et al., 2007). Shortening of the poly(A) tail is generally mediated by the CCR4-NOT complex (Yamashita et al., 2005). To test whether the CCR4-NOT complex is required for mRNA decay during the M-G1 phase transition, we depleted CNOT1, the scaffold subunit of the CCR4-NOT complex, using siRNA-mediated depletion in RPE-1 cells (Fig. S4A). For initial experiments, we focused on TOP2A and CDK1 mRNAs, as both were rapidly and robustly degraded at the M-G1 phase transition. In line with the fact that CNOT1 is essential for proliferation (Blomen et al., 2015; Hart et al., 2015; Wang et al., 2015), fewer mitotic cells were present in the cultures following CNOT1 depletion. Nonetheless, in the mitotic cells that could be identified, depletion of CNOT1 caused a 20–25% increase in the relative abundance of both TOP2A and CDK1 mRNAs in telophase (when decay of these mRNAs has normally occurred) compared to control cells (Fig. 4A). These results suggest that CNOT1-dependent mRNA deadenylation is important for mRNA decay at the M-G1 phase transition.

**Figure 4.**
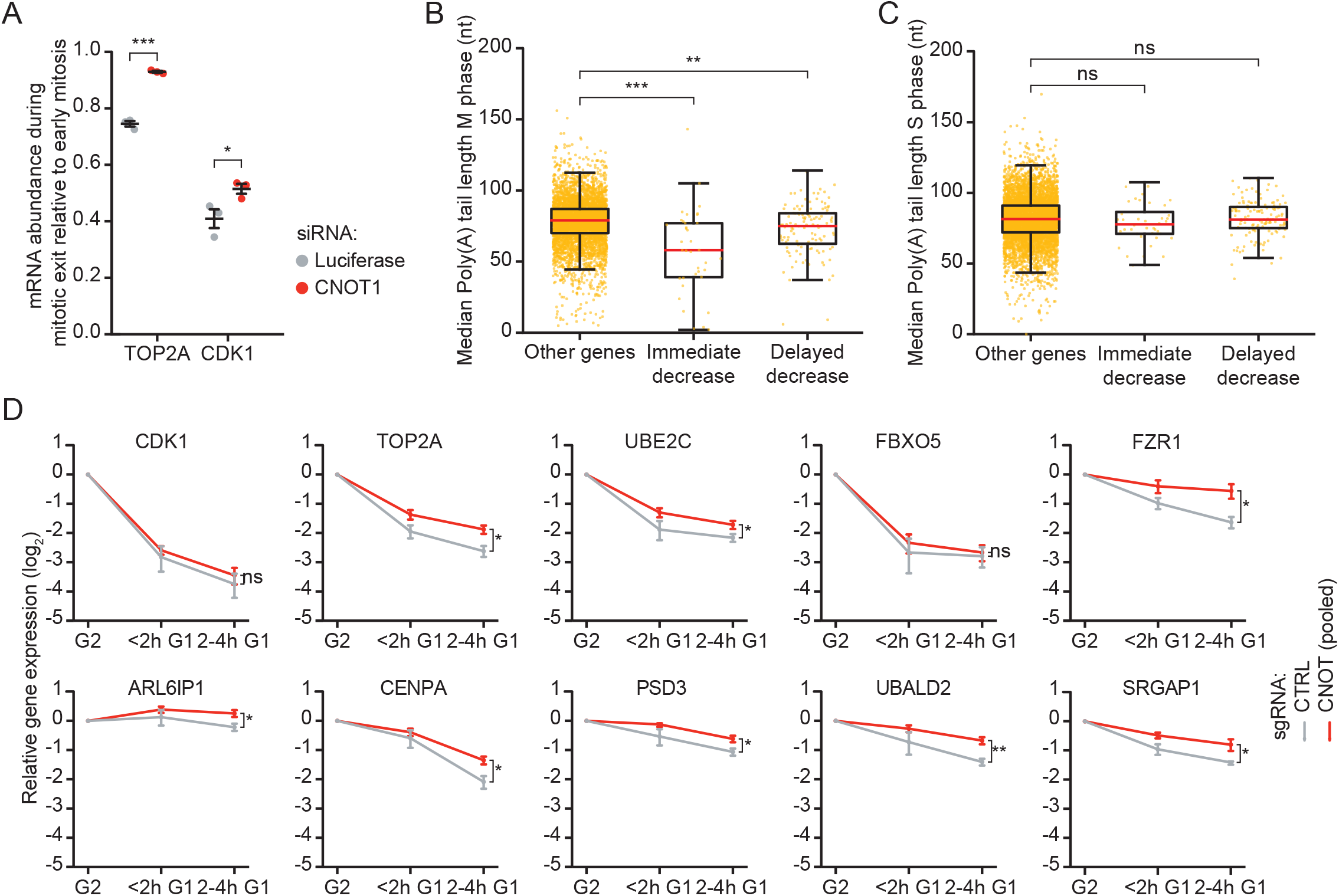
CNOT1 promotes decay of mRNAs during mitotic exit and early G1 phase. **A)** Cells were transfected with indicated siRNAs. Two days after transfection, cells were fixed and TOP2A and CDK1 mRNAs were stained using smFISH. The mRNA levels were quantified using ImageJ. To calculate the relative abundance of mRNAs during mitotic exit, we divided the number of mRNAs present in telophase by the average number of mRNAs present in prophase, prometaphase and metaphase (mRNA abundance is similar during these phases of mitosis (Fig. 2D-F). Note that relative abundance was used instead of absolute copy number, as the absolute number of identifiable foci varied between experiments due to variations in labeling intensity of smFISH probes. Each dot represents a single experiment and lines with error bars indicate mean ± SEM of three independent experiments. Per experiment, at least 10 cells during mitotic exit and 10 early mitotic cells were quantified (See Supplementary table 2). P-values are based on a one-tailed Student’s t-test. **B-C)** Boxplot of poly(A)-tail lengths in mitosis (B) and S phase (C) (from Park et al., 2016) for genes subject to decay during mitotic exit (*Immediate decrease*), during early G1 phase (*Delayed decrease*), or genes that are not subject to decay (Other genes). P-values are based on a one-tailed Student’s t-test. **D)** CNOT1 contributes to mRNA decay at the M-G1 phase transition for many genes. RPE-FUCCI CRISPRi cells, infected with control- or CNOT1-targeting sgRNAs, were sorted into populations of G2/M phase and G1 phase cells at 5 days post sgRNA infection. The mRNA levels of indicated genes were measured by RT-qPCR. Per experiment, mRNA expression was measured in three CNOT1-depleted samples (using 3 independent sgRNAs targeting CNOT1) and one control sample. mRNA levels of the 3 independent samples were averaged for each biological replicate. Lines and error bars indicate average ± SEM of 3 individual experiments. P-values are based on a one-tailed Welch’s t-test. P-values are indicated as * (p < 0.05), ** (p < 0.01), *** (p < 0.001), ns = not significant.

A previous study found that the mRNAs of many cell cycle genes contain significantly shorter poly(A) tails in M phase compared to S phase (Park et al., 2016). Interestingly, re-analysis of their data revealed that the poly(A)-tails of transcripts in the *immediate decrease* group were shorter than those in the control group (i.e. genes that did not show mRNA decay at the M-G1 phase transition) (median value 58 versus 79, respectively) (Fig. 4B). The *delayed decrease* group mRNAs also have shorter poly(A) tails than those in the control group during mitosis, although the effect was relatively minor (median value 75 versus 79, respectively). Importantly, the shortened poly(A) tails were specific to mitosis, as poly(A) tail lengths in S phase of *immediate* and *delayed decrease* groups were similar to those of control genes (Fig. 4C). Collectively, these data show that CNOT1 is important for the decay of TOP2A and CDK1 mRNAs during the M-G1 phase transition and suggest that CNOT1-dependent deadenylation in mitosis may contribute to the decay of many mRNAs at the M-G1 phase transition.

To determine whether CNOT1 is also involved in the second wave of mRNA decay, we depleted RPE-FUCCI cells of CNOT1 using CRISPR interference (CRISPRi) (Gilbert et al., 2013, 2014). Using CRISPRi, we could knock down CNOT1 in a large population of cells, allowing subsequent FACS-based isolation of sufficient numbers of early G1 phase cells. We used three independent single guide (sg)RNAs to target CNOT1 by CRISPRi, which resulted in a modest (∼50%) reduction of CNOT1 mRNA levels (Fig. S4B). Nonetheless, the modest reduction of CNOT1 mRNA levels causes a clear cell cycle arrest (Fig. S4C), confirming the essential function of CNOT1 for normal proliferation. Importantly, CNOT1 depletion did not affect the synthesis rates of the FUCCI-G1 fluorescent reporter (i.e. accumulation of fluorescence over time) during G1 phase (Fig. S4D), allowing us to isolate control and CNOT1-depleted cells at similar times in G1 phase based on FUCCI fluorescence through FACS sorting. G2/M phase and early G1 phase cell populations were isolated by FACS and mRNAs of the *immediate decrease* and *delayed decrease* groups were measured by RT-qPCR. Even though depletion of CNOT1 was modest in these experiments, a small but reproducible decrease in decay was observed for eight of the ten genes tested (Fig. 4D). Taken together, these data demonstrate that both waves of post-mitotic mRNA decay are stimulated by CNOT1.

## Discussion

### Assigning a precise cell cycle time to individual, sequenced cells

Understanding of the regulation and heterogeneity of gene expression has flourished due to the development of single cell sequencing techniques. To investigate transcriptome changes over time, trajectory inference methods have been developed that allow *in silico* ordering of cells, based on (dis)similarities in transcriptomes (Saelens et al., 2019). This creates single cell trajectories of a biological process of interest, such as differentiation or the cell cycle (Fig. S1J) (Haghverdi et al., 2016; Trapnell et al., 2014), and is useful to study dynamics in gene expression. However, due to clustering based on transcriptome (dis)similarities, pseudo time may under/overestimate true cellular state durations (Tian et al., 2019). In addition, trajectories lack real temporal information and are therefore not ideal to determine absolute mRNA synthesis and degradation rates. To circumvent these issues, we have developed a method that combines live-cell microscopy and FACS-analysis of the FUCCI system with scRNA-seq, to generate a high-resolution, time-resolved transcriptome profile of the M-G1 phase transition in human cells. Even though the FUCCI system has previously been used to order single cell transcriptomes along the cell cycle (Battich et al., 2020; Hsiao et al., 2020; Mahdessian et al., 2021), a unique feature of our method is that it uses precisely calibrated FUCCI reporter fluorescence intensities for accurate assignment of cell cycle times of individual, sequenced cells. We use these calibrated fluorescence intensities to align cells along the cell cycle according to their cell cycle ‘age’. We identify hundreds of mRNAs that show sharp transitions in their expression levels as cells progress from mitosis to G1 phase. The availability of temporal information allowed us to quantitatively determine mRNA degradation rates for these transcripts, which is not possible using trajectory inference methods. Similar approaches are likely possible for other biological events for which fluorescence reporters are available, making this approach a broadly applicable method.

### mRNA decay at the M-G1 phase transition ‘resets’ the transcriptome

It is evident that the expression of G2/M-specific genes is reduced following the completion of cell division through scheduled protein degradation and transcriptional inactivation (Bar-Joseph et al., 2008; Castro et al., 2005; Chaudhry et al., 2002; Cho et al., 1998, 2001; Grant et al., 2013; Harper, 2002; Nakayama and Nakayama, 2006; Peters, 2002; Vodermaier, 2004; Whitfield et al., 2002). Here, we identify widespread scheduled mRNA decay during mitotic exit and early G1 phase as an additional mechanism acting to reset gene expression following cell division. Why would mRNA transcripts be actively degraded when the clearance of transcripts will eventually be achieved by transcription shut-down alone? All the mRNAs we tested are stable during mitosis and late G1 phase (Fig. 3D-E). Therefore, transcription inhibition by itself would lead to significant carry-over of transcripts into the next cell cycle. Considering the half-live of these mRNAs in G2 and M phase, they would persist in G1 phase for many hours. Therefore, decay-mediated clearance of mRNAs as cells exit mitosis will aid to limit expression of the encoded proteins in G1 phase, especially since the majority of degraded mRNAs is efficiently translated during G1 phase (Tanenbaum et al., 2015). As these genes include many genes that encode for proteins with important functions in cell cycle control (Fig. S1I), their continued expression in G1 phase may perturb normal cell cycle progression, and could potentially even contribute to cellular transformation (García-Higuera et al., 2008; Park et al., 2008; Sigl et al., 2009). Thus, scheduled mRNA decay during the cell cycle may be important to restrict gene expression of many cell cycle genes to their appropriate cell cycle phases.

### CNOT1 promotes two waves of mRNA decay during the M-G1 phase transition

We have identified two consecutive waves of mRNA decay as cells progress through mitosis and into G1 phase (Fig. 2A,C and S3G). The fact that mRNA decay occurs in two waves may indicate the existence of two distinct mechanisms that act consecutively to degrade transcripts. Interestingly, these two waves of mRNA degradation during the M-G1 phase transition are highly reminiscent of the two consecutive waves of protein degradation that occur at the same time (Alfieri et al., 2017; Sivakumar and Gorbsky, 2015).

The regulation of mRNA decay often occurs through (sequence) specific interactions between mRNAs and RNA binding proteins (RBPs). Through direct interactions with mRNAs, RBPs can recruit components of the RNA decay machinery, such as the CCR4-NOT complex, to the mRNA. Recruitment of CCR4-NOT, a key regulator of gene expression, will then induce deadenylation of the target transcript, generally followed by degradation (Garneau et al., 2007). Interestingly, a previous report found that the poly(A)-tail lengths of the genes we identified as *immediate decay* are already shortened in early mitosis, before these mRNAs are degraded (Fig. 4B-C) (Park et al., 2016). The observation that poly(A) tails of transcripts decayed during the M-G1 phase transition are shorter in mitosis could suggest that CCR4-NOT-dependent deadenylation during early mitosis marks these transcripts for subsequent decay during mitotic exit and early G1 phase. Indeed, we identified CNOT1, an essential member of the CCR4-NOT complex, as a regulator of post-mitotic mRNA decay (Fig. 4A and 4D). We note that the effects of CCR-NOT depletion on mRNA decay are modest in our experiments, but the magnitude of the effect is likely caused, at least in part, by the inability to effectively deplete CNOT1, while maintaining cells in a cycling state. Perhaps rapid degradation of CNOT1 protein using inducible protein degradation systems could solve this issue in the future (Banaszynski et al., 2006; Nishimura et al., 2009; Yesbolatova et al., 2020).

Our data shows that both waves of mRNA decay during the M-G1 phase transition are stimulated by CNOT1 (Fig. 4D). Nonetheless, these waves of mRNA decay may be regulated independently, involving distinct RBPs. Binding of distinct RBPs could ensure the timely decay of specific sets of mRNA during either mitotic exit or early G1 phase. It will be interesting to investigate which RBPs are involved in recognizing different subsets of mRNAs that need to be degraded during particular times in the cell cycle. Identification of such RBPs will allow a better understanding of the function and mechanisms of scheduled mRNA degradation during the cell cycle.

## Acknowledgements

We thank members of the Tanenbaum group for helpful discussions, and Xiaowei Yan for help during the initial stages of the project. We thank the Hubrecht Institute flow cytometry facility and single cell sequencing facility (now Single Cell Discoveries) for their technical support. This work was financially supported by the European Research Council (ERC) through an ERC starting grant (ERCSTG 677936-RNAREG) to M.E.T.; M.E.T. is also supported by the Oncode Institute that is partially funded by the Dutch Cancer Society (KWF).

## Author contributions

L.K. and M.E.T. conceived the project. L.K. and S.S. performed the experiments and analysis, and S.S. performed the computational modeling. L.K. and S.S. prepared the figures, and L.K., S.S, and M.E.T. wrote the manuscript.

## Figure legends

**Figure S1 – Supplementary to figure 1**.

**A)** Fluorescence microscopy time traces of RPE-1 cells expressing FUCCI-G1 and FUCCI-G2 markers. Asynchronously growing RPE-FUCCI cells were imaged every 5 minutes and the average nuclear intensities for both FUCCI markers was measured using Image J (red and green lines, respectively). Dark red and green lines represent the average of 30 cells from one experiment, light red and green lines represent individual cells.

**B)** Microscopy-based analysis of fluorescence intensities of FUCCI-G1 marker in RPE-FUCCI cells. These data were used to identify the average FUCCI-G1 fluorescence level in early S phase cells (yellow dots, see methods for details).

**C)** FACS plots of FUCCI fluorescence after different durations of Taxol treatment. To quantify the increase in FUCCI-G1 fluorescence over time, asynchronously growing RPE-FUCCI cells were treated with Taxol for the indicated durations to arrest cells in mitosis, thus preventing new cells from entering G1 phase. Over time, FUCCI-G1 fluorescence increases, which results in the gradual loss of cells with low FUCCI-G1 fluorescence. The lowest FUCCI-G1 fluorescence intensity after a 1-, 2- or 4-hour incubation with Taxol were identified (dotted lines), and used to calculate the FUCCI-G1 fluorescence intensity relative to early S phase (see methods).

**D)** Comparison of the relative FUCCI-G1 fluorescence intensities as determined by the polynomial equation and by Taxol treatment of cells followed by FACS (see panel D) at 1, 2 and 4 hours after mitosis. Error bars indicate SEM of 3 experiments

**E-G)** Comparison of normalized read counts from three sequencing experiments. Each dot represents the average G2 level of a gene. Dotted red line indicates similar average read counts in both plates.

**H)** Histogram showing the position in the cell cycle of all cells subjected to scRNA-Seq.

**I)** Gene Ontology of genes downregulated in G1 phase compared to G2/M phase, identified when positioning cells along the cell cycle based on FUCCI marker fluorescence.

**J)** Single-cell trajectory of the M-G1 phase transition constructed by Monocle2. Colors indicate the cell cycle position based on FUCCI marker fluorescence.

**K)** Differential transcriptome analysis of G1 vs G2/M phase RPE-FUCCI cells, aligned based on Monocle2 trajectory inference.

**Figure S2 – Supplementary to figure 2**.

FACS analysis of asynchronously growing RPE-FUCCI cells, including gating strategy for the identification of various populations of G1, early S and G2 phase cells (left). The table displays the FUCCI-G1 fluorescence intensity relative to early S phase (middle column) that is associated with the time a cell has spent in G1 phase (left column), based on the polynomial equation (Fig. 1B). The relative FUCCI-G1 fluorescence intensity values (middle column) are converted to absolute FUCCI-G1 fluorescence intensity values (right column) using the mean FUCCI-G1 fluorescence intensity during early S phase (value listed above the table). The absolute fluorescence intensities of FUCCI-G1 (right column) were then used to isolate cells of a specific age with the corresponding gating strategy shown in the FACS-plot on the left.

**Figure S3 – Supplementary to figure 3**

**A-F)** mRNA abundance over time of genes subject to post-mitotic mRNA decay. Blue lines indicate best fit obtained using the mathematical model described in Fig. 3A. Example genes are shown that are representative for the *immediate decrease* group (CDK1, TOP2A and UBE2C) and for the *delayed decrease* group (CENPA, ALR6IP1 and UBALD2).

**G)** mRNA levels over time were fit as in (A-F) and the onset time of the decline in mRNA levels was determined for each of the 220 downregulated genes.

**H-J)** Comparison of mRNA half-lives during the M-G1 phase transition as calculated in Fig. 3B with mRNA half-lives in asynchronous cells determined in previous studies (Herzog et al., 2017; Schwanhäusser et al., 2011; Tani and Akimitsu, 2012). Dashed lines indicate identical half-lives. Note that the half-lives of most genes are shorter during the M-G1 phase transition than in asynchronous growing cells.

**K)** Boxplot of mRNA half-lives of *immediate* and *delayed decrease* genes. For each gene, the half-live was determined from the moment mRNA levels start to decrease (see Supplementary table 1). P-value is based on a one-tailed Student’s t-test.

**L)** Analysis of transcription inhibition by Actinomycin D. Expression levels of the DNA damage-induced gene CDKN1a were measured by RT-qPCR in cells that were DNA damaged (exposed to 5 Gy ionizing radiation), in the presence or absence of Actinomycin D, relative to non-irradiated cells. Each dot represents a single experiment and lines with error bars indicate mean ± SEM.

**M)** Mitotic index of RPE-1 cells treated with the transcription inhibitor Actinomycin D. RPE-1 cells were arrested in G2 using a CDK1 inhibitor (RO 3306). After 16h, the CDK1 inhibitor was removed and replaced by Taxol, thereby blocking cells in mitosis. 45 minutes later, mitotic cells were collected through mitotic shake-off, after which Actinomycin D was added for up to 2 hours. Cells were fixed and the fraction of mitotic cells was determined by FACS (by staining cells for DNA content and the mitosis-specific marker phosphorylated histone 3 at ser 10). Each dot represents a single experiment and lines with error bars indicate mean ± SEM. P-values are indicated as * (p < 0.05), ** (p < 0.01), *** (p < 0.001), ns = not significant.

**Figure S4 – Supplementary to figure 4**

**A)** Validation of siRNA-mediated knockdown. RPE-1 cells were transfected with indicated siRNAs, or a control siRNA targeting luciferase. mRNA levels relative to control were measured by RT-qPCR at 48 hours post siRNA transfection. Lines with error bars indicate the average ± SEM of three independent experiments.

**B)** CNOT1-depletion levels of cells depicted in Fig. 4D. RPE-FUCCI CRISPRi cells were infected with the indicated sgRNAs. 5 days post infection, cells were sorted into the indicated cell cycle fractions, and CNOT1 expression was measured by RT-qPCR. Error bars indicate the SEM of three individual experiments.

**C)** Cell cycle distribution of control and CNOT1-depleted RPE-FUCCI cells. Cells were treated as in Fig. 4B, except their cell cycle distribution was determined by FACS, based on FUCCI fluorescence. Error bars represent SEM of three independent experiments.

**D)** Accumulation of the FUCCI-G1 marker in control or CNOT1-depleted RPE-FUCCI cells. RPE-FUCCI CRISPRi cells were infected with the indicated sgRNAs. 5 days post infection, the cells were imaged using time-lapse microscopy, and FUCCI-G1 fluorescence was determined for cells at metaphase and 2 or 4 hours thereafter. Fluorescence intensities were normalized against the average fluorescence intensity of control cells at 2 hours post-metaphase. Lines and error bars indicate the average ± SEM of three independent experiments. At least 8 cells per condition per experiment were quantified (see Supplementary table 2). P-values are based on a one-tailed Student’s t-test. P-values are indicated as * (p<0.05), ** (p<0.01), *** (p<0.001), ns = not significant.

## Methods

### Transfections and lentivirus production

Lentivirus was produced by transfecting HEK293T cells with packaging plasmids (pMD2.G and psPAX2; addgene #12259 and #12260, respectively) and lentiviral plasmids carrying the transgene of interest. Plasmids were transfected using FuGENE HD (Promega) according to the manufacturer’s protocol. Two days post transfection, virus was harvested by collecting the culture medium, pelleting cell debris by centrifugation, and collecting the supernatant.

### Cell culture and generation of cell lines

HEK293T cells were maintained in Dulbecco’s Modified Eagle Medium (DMEM) supplemented with 5% fetal bovine serum (FBS, Sigma-Aldrich) and 1% penicillin/streptomycin (Gibco). RPE-1 cells and derivatives were maintained in DMEM/Nutrient Mixture F-12 (DMEM/F12, Gibco) supplemented with 10% FBS and 1% penicillin/streptomycin. To generate RPE-FUCCI cells, RPE-1 cells were transduced with lentivirus expressing mkO2-hCdt1(30/120) (FUCCI-G1) and lentivirus expressing mAG-hGem(1/110) (FUCCI-G2) (Sakaue-Sawano et al., 2008). Single clones were isolated by fluorescent activated cell sorting (FACS) on a BD FACSFUSION system. One clone was selected that showed cyclic expression of both reporter constructs. To generate RPE-FUCCI CRISPRi cells, RPE-FUCCI cells were transduced with lentivirus carrying dCas9-BFP-KRAB (Jost et al., 2017), and the 15% highest BFP-positive cells were isolated by FACS.

*Synchronization of cells in mitosis, transcription inhibition and FACS analysis of mitotic cells* In order to synchronize cells in mitosis, we first arrested cells in G2 by treating cells with the CDK1-inhibtor RO-3306 for 16 hours. Subsequently, RO-3306 was removed and the cells were washed twice with PBS before applying fresh medium supplemented with Taxol, which blocks cells in mitosis. Finally, 45 minutes after Taxol addition, mitotic cells were separated from the interphase cells by shaking of the culture dish (shake-off). This specifically detaches mitotic cells, that were then harvested by collecting the culture medium. To inhibit transcription, we treated cells with 1µg/ml Actinomycin D (Sigma-Aldrich) for the indicated durations. In order to identify mitotic cells, cells were fixed in 80% ethanol (−20°C). Thereafter, cells were stained using an antibody targeting the mitosis-specific marker phosphorylated histone 3 (4N pH3)-ser10 (Upstate, 06-570) and propidium iodide to label DNA content. The mitotic fraction was determined as the fraction of 4N pH3-ser10 positive cells.

### siRNA transfections

Cells were grown in 96-well microscopy plates (Matriplate, Brooks) or 96-well culturing plate (Greiner Bio-one) and siRNAs were transfected at a final concentration of 10 nM using RNAiMAX (Invitrogen) according to the manufacturer’s protocol. Two days post-transfection, cells were either fixed for smFISH or the RNA was harvested for RT-qPCR analysis. For knockdown of CNOT1 and DCP2 we used ON-TARGET plus siRNAs from Dharmacon. As a control, we used a custom siRNA targeting luciferase (5’-CGUACGCGGAAUACUUCGAUU-3’) from Dharmacon.

### CRISPRi

For CRISPR interference (CRISPRi), RPE-FUCCI CRISPRi cells were infected with lentivirus particles expressing a non-targeting single-guide RNA (sgRNA), or a sgRNA targeting CNOT1 (Horlbeck et al., 2016) and a puromycin resistance cassette followed by BFP. Two days post infection, infected cells were selected with puromycin for 3 days to eliminate uninfected cells. Sequences of sgRNAs used in this study can be found in Supplementary table 3.

### Quantitative reverse transcription PCR (RT-qPCR)

For RT-qPCR analysis, cells were lysed in TriSure (Bioline) and RNA was extracted according to the manufacturers’ protocol. First strand synthesis was performed using Bioscript (Bioline). mRNA expression levels were quantified using SYBR-Green Supermix (Bio-Rad) on a Bio-Rad Real-time PCR machine (CFX Connect Real-Time PCR Detection System). Relative mRNA expression levels were calculated using the ΔΔCt-method. GAPDH and RPN1 were selected as reference genes for normalization, based on their reported high mRNA stability (Schwanhäusser et al., 2011). RT-qPCR primers were designed using Primer3, for sequences see Supplementary table 3.

### Branched DNA single molecule Fluorescent In Situ Hybridization

Single molecule fluorescent *in situ* hybridization (smFISH) was performed using viewRNA probes targeting TOP2A (probe# VA1-14609) and CDK1 (probe# VA6-17545) (ThermoFisher). Staining was done according to the manufacturer’s protocol. In brief, cells were grown in 96-well microscopy plates (Matriplate, Brooks) and fixed for 30 minutes using 4% formaldehyde. Then, cells were permeabilized with detergent solution for 5 minutes at room temperature (RT), and subsequently treated with protease solution for 10 minutes at RT. To label the RNAs, cells were incubated with probes targeting TOP2A and CDK1 for 3 hours at 40°C. Subsequent probes (preAmplifier, Amplifier and Label Probe) were incubated for 1 hour at 40°C. Between probe incubations, cells were washed with wash buffer for 3x 1 minute. After the final incubation (with Label Probe), cells were washed and incubated with DAPI (ThermoFischer, D1306) and wheat germ agglutinin, conjugated to Alexa Fluor 488 (ThermoFisher, W11261) to label DNA and membranes, respectively.

### Microscopy

For live-cell microscopy, RPE-FUCCI cells were grown on microscopy plates and imaged using a Nikon Ti-E with PFS, equipped with an Andor Zyla 4.2Mpx sCMOS camera, CFI S Plan Fluor ELWD 20x air objective (0.45 NA) and a Lumencor SpectraX light source. Temperature and CO_2_ control were provided by an OKO-lab Boldline microscope cage and CO_2_ controller. Image analysis was performed using ImageJ software.

For imaging of bDNA-FISH stained samples, we used a Nikon TI2 inverted microscope with a perfect focus system, equipped with a Yokagawa CSU-X1 spinning disc, a 100x oil objective (1.49 NA), and a Prime 95B sCMOS camera (Photometrics).

### SORT-Seq

SORT-Seq was performed as described previously (Muraro et al., 2016). Briefly, we sorted in total 104 G2/M phase cells (FUCCI-G1 negative and FUCCI-G2 positive cells, Fig. 1C) and 893 G1 phase cells (FUCCI-G1 positive and FUCCI-G2 negative cells, Fig. 1C) in three 384-wells plates. Each 384-wells plate contained G1 phase cells from 0-4 hours after the start of G1 phase, but only one plate contained G1 phase cells from 4-9 hours after the start of G1 phase. Therefore, we only used cells from 0-4 hours after the start of G1 phase to identify differentially expressed genes. In subsequent analyses (i.e. the spline analysis and the modelling) we did use all G1 phase cells. After sequencing, we continued with cells (841 in total) that passed quality tests (we removed cells with less than 5900 UMIs or more than 111.000 UMIs to lose low quality cells and doublets, respectively). Finally, we normalized for differences in mRNA recovery per cell using Monocle2 (R package).

### Cell cycle timing using the FUCCI system

To obtain a temporal transcriptome profile as cells progress from mitosis into G1 phase, we wanted to compute a cell cycle time for each sorted G1 phase cell (that is, how much time a cell has spent in G1 phase at the moment of sorting). Since FUCCI-G1 levels positively correlate with the amount of time a cell has spent in G1 phase, we reasoned that we could use the measured FUCCI-G1 levels to infer a cell cycle time for a G1 phase cell. To characterize precisely how FUCCI-G1 levels increase during G1 phase, we imaged RPE-FUCCI cells under the microscope with a time-interval of 5 minutes and selected cells that progressed through mitosis into G1 phase. Next, we measured the mean fluorescence intensities of both FUCCI sensors in a region of interest (ROI) in the nucleus using ImageJ and subtracted background signal measured in an extracellular ROI. In each experiment we quantified the fluorescence intensities of the FUCCI sensors in 30 cells.

To compute the average FUCCI-G1 time-trace during G1 phase, we aligned the time-traces of individual cells at the metaphase-to-anaphase transition, which is defined by a sudden decrease in FUCCI-G2 fluorescence (Sakaue-Sawano et al., 2008). Next, since the total amount of time a cell spends in G1 phase differs for each cell, we clipped individual time-traces at the end of G1 phase, which ends shortly after the FUCCI-G2 levels start to increase (Grant et al., 2018). To determine the moment the FUCCI-G2 levels start to increase, we first corrected the FUCCI-G2 traces for fluorescence crosstalk from the FUCCI-G1 marker, which is also excited by the 488 nm laser used for imaging of the FUCCI-G2 marker. To correct the FUCCI-G2 time-traces for crosstalk from the FUCCI-G1 marker, we subtracted at each time point 31% of the FUCCI-G1 fluorescence intensity from the FUCCI-G2 fluorescence intensity. Next, we determined the time point when the mean FUCCI-G2 fluorescence intensity reached a threshold value, which was set by visual inspection, and clipped all FUCCI-G1 time traces at the time point of FUCCI-G2 increase. Finally, we computed for each experiment the average FUCCI-G1 levels from the moment of metaphase and fit the average of three experiments to a third-order polynomial (Fig. 1B).

To directly compare the FUCCI-G1 levels that we measured on the microscope to the FUCCI-G1 levels that we measured on the FACS, we normalized both microscopy- and FACS-measured FUCCI-G1 levels to the average FUCCI-G1 level of early S phase cells. To quantify the mean FUCCI-G1 fluorescence intensity in early S phase cells on the microscope, we analyzed the mean nuclear intensities of at least 700 cells per experiment (Fig. S1B). As above, we compensated for fluorescence crosstalk from the FUCCI-G1 marker into the FUCCI-G2 channel by subtracting 31% of the FUCCI-G1 fluorescence intensity from the FUCCI-G2 fluorescence intensity. Next, we determined the range of fluorescence intensities for both the FUCCI-G1 and FUCCI-G2 markers (by subtracting the lowest fluorescence intensity from the highest fluorescence intensity), and defined the early S phase population as those cells with FUCCI-G2 intensities between 2,5% and 10% of the range of FUCCI-G2 intensities and FUCCI-G1 intensities higher than 2,5% of the range of FUCCI-G1 intensities (Fig. S1B, yellow dots). We computed the average FUCCI-G1 level of the early S phase cells, and normalized the third-order polynomial fit against the average FUCCI-G1 level of the early S phase cells.

To quantify the mean FUCCI-G1 fluorescence intensity in early S phase cells on FACS, we analyzed the FUCCI sensors on FACS (Fig. 1C). We define the early S phase population as those cells that have high FUCCI-G1 levels and have started to increase the expression of the FUCCI-G2 marker Fig. 1C), and computed the average FUCCI-G1 level in early S phase cells. To obtain a cell cycle time for each G1 phase cell that was sequenced, we determined the FUCCI-G1 fluorescence intensity level that was obtained by FACS and normalized the FUCCI-G1 level to the average early S phase FUCCI-G1 value. Finally, we used the third-order polynomial fit to infer the cell cycle time of each G1 phase cell from its normalized FUCCI-G1 level.

### Cell cycle timing using Monocle2

To rank cells using Monocle2 (R package), we used all G2/M phase cells and G1 phase cells that were from the first 4 hours of G1 phase (based on FUCCI cell cycle timing; see section *‘SORT-seq’*). Next, we performed an initial differential transcriptome analysis comparing G2/M phase (FUCCI-G1 marker negative and FUCCI-G2 marker positive) and G1 phase cells (FUCCI-G1 marker positive and FUCCI-G2 marker negative) to select differentially expressed genes that Monocle2 can use in subsequent steps to reconstruct the single-cell trajectory. Monocle2 selected a total of 430 genes that were used to reconstruct the single-cell trajectory, and both the cell rank and the Monocle2 assigned pseudo-times were compared to FUCCI-based ranking and cell cycle timing.

### Differential transcriptome analysis

Differential transcriptome analysis was performed with Monocle2 (R package), either using FUCCI-based or Monocle2-based cell cycle time. For the differential transcriptome analysis we used all G2/M phase cells and only G1 phase cells from the first 4 hours of G1 phase (see section *‘SORT-seq’*). To increase the confidence of our differential transcriptome analysis, we only selected genes for analysis that were clearly detected in all three 384-wells plates. To select detected genes, we computed for each gene in each single 384-wells plate its average expression in G2/M phase cells (as we didn’t want to bias against genes that were downregulated in G1 phase), and only selected genes which had an average expression of at least 2 reads in each single 384-wells plate. This resulted in a dataset of 3985 genes. Finally, after differential transcriptome analysis, genes that showed at least a twofold increase or decrease in expression and had at least a p-value of 1E^-5^ (based on a Bonferroni correction from a p-value of 0.05) were selected as upregulated or downregulated genes, respectively.

### Spline analysis

For the spline analysis (performed in Matlab R2018b), we used the full set of 841 cells (see section ‘*SORT-seq’*). We selected the 220 genes that were identified in the differential transcriptome analysis as downregulated in G1 phase, and fit each gene profile with a smoothing spline. Next, we computed the derivative of the splines at each time point and determined the time when the derivative was minimal for each gene (i.e. the moment mRNA levels decreased most). To compare different genes to each other, we normalized the derivative of each gene to its minimum value (i.e. setting the minimum value to −1). Finally, we determined for each gene the first time point during which the normalized derivative was at least −0.95 (where −1.0 is the minimum slope after normalization), and divided genes in two groups; one group in which the minimum slope was reached at the first time point (i.e. during mitosis) and one group in which the minimum slope was reached during G1.

### Modelling mRNA decrease

mRNA levels (*m*) depend on the synthesis (*µ*) and degradation (γ) rate, and the change in mRNA levels over time can be described as follows.

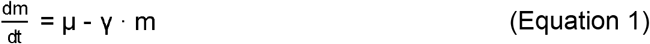

To describe the mRNA levels as cells progress from mitosis into G1 phase, we assumed a simple model in which the observed decrease of mRNA levels is explained by a decrease in the synthesis rate and/or an increase in the degradation rate at a specific time point during M or early G1 phase (referred to as the onset time or *t*_*onset*_). When mRNA levels start at a given value (*m*_*0*_), the solution of equation 1 results in the following expression for the mRNA levels over time.

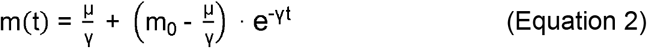

Furthermore, we assumed that mRNA levels remain constant before the onset time, resulting in the following pair of equations to describe the mRNA levels as cells progress from mitosis into G1 phase.

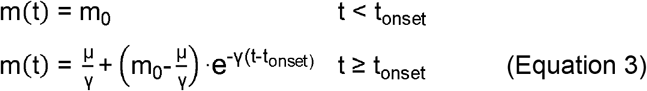

For each gene, we optimized *t*_*onset*_ (performed in Matlab R2018B) using an iterative search (between 0 and 370 minutes after metaphase in steps of 10 minutes), in which we optimized *m*_*0*_, *µ*, and γ using least square fitting for each *t*_*onset*_. Finally, we computed a sum of squared errors (SSE) between the data (using the full dataset of 841 cells) and model for each *t*_*onset*_ and selected the *t*_*onset*_ with the minimal SSE.

### Calculating half-lives

We computed mRNA half-lives from the degradation rates (γ) (that we obtained from the modelling) using equation 4.

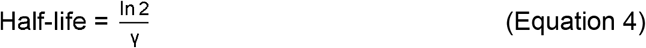

### Statistics

Statistical comparisons were made using a unpaired one-tailed Student’s t-test (Fig. 2E, 2F, 3D, 3E, 4A, 4B, 4C, S3K and S4D), a paired one-tailed Student’s t-test (Fig.3C) or a one-tailed Welch’s t-test (Fig. 4D)

