## Supplementary figures and images for "Time-resolved single-cell sequencing identifies multiple waves of mRNA decay during mitotic exit"

### Supplemental Figure 1

Figure S1 - Supplemental to Figure 1

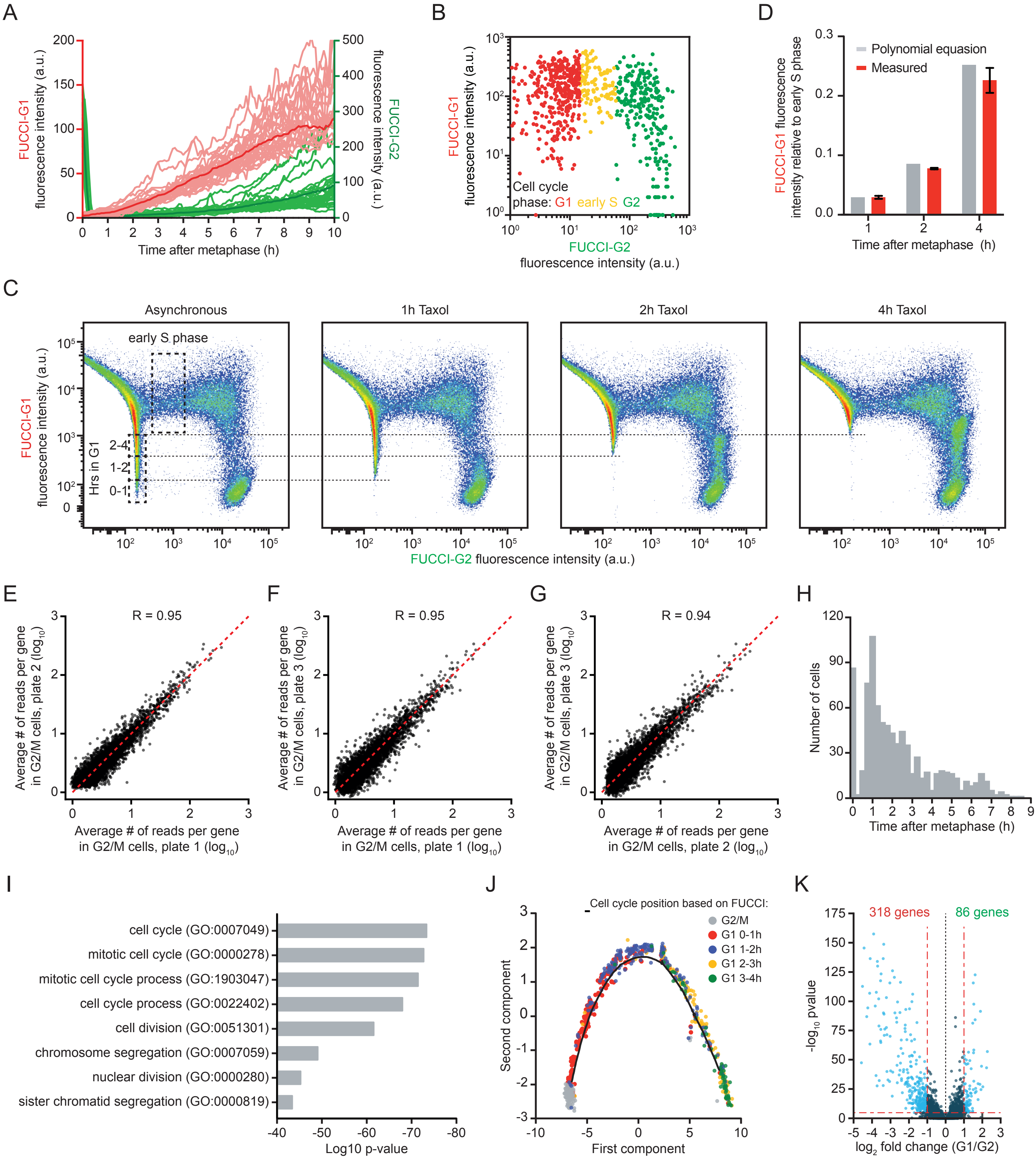

### Supplemental Figure 3

Figure S3 - Supplementary to Figure 3.

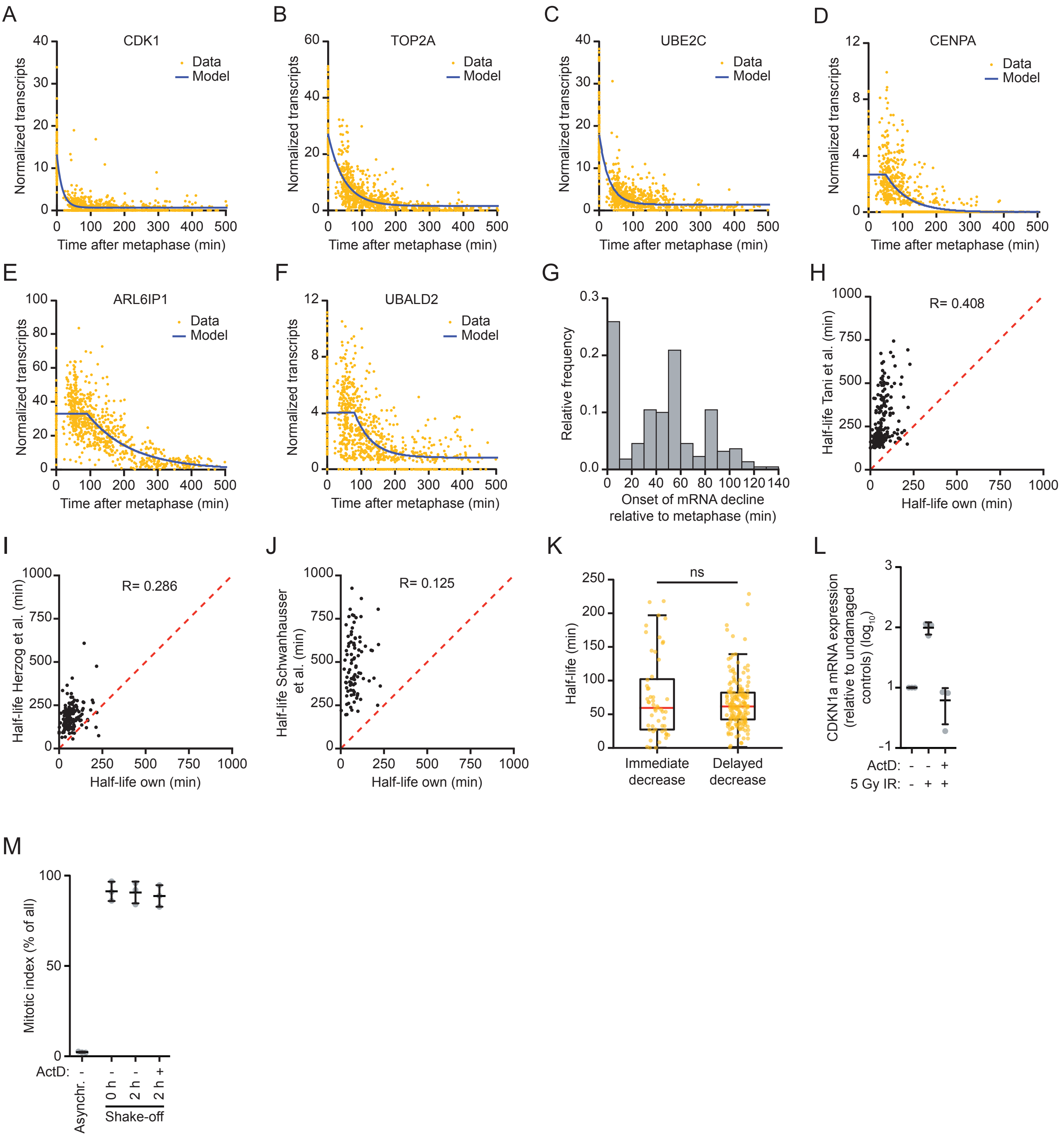

### Supplemental Figure 4

Figure S4 - Supplementary to Figure 4.

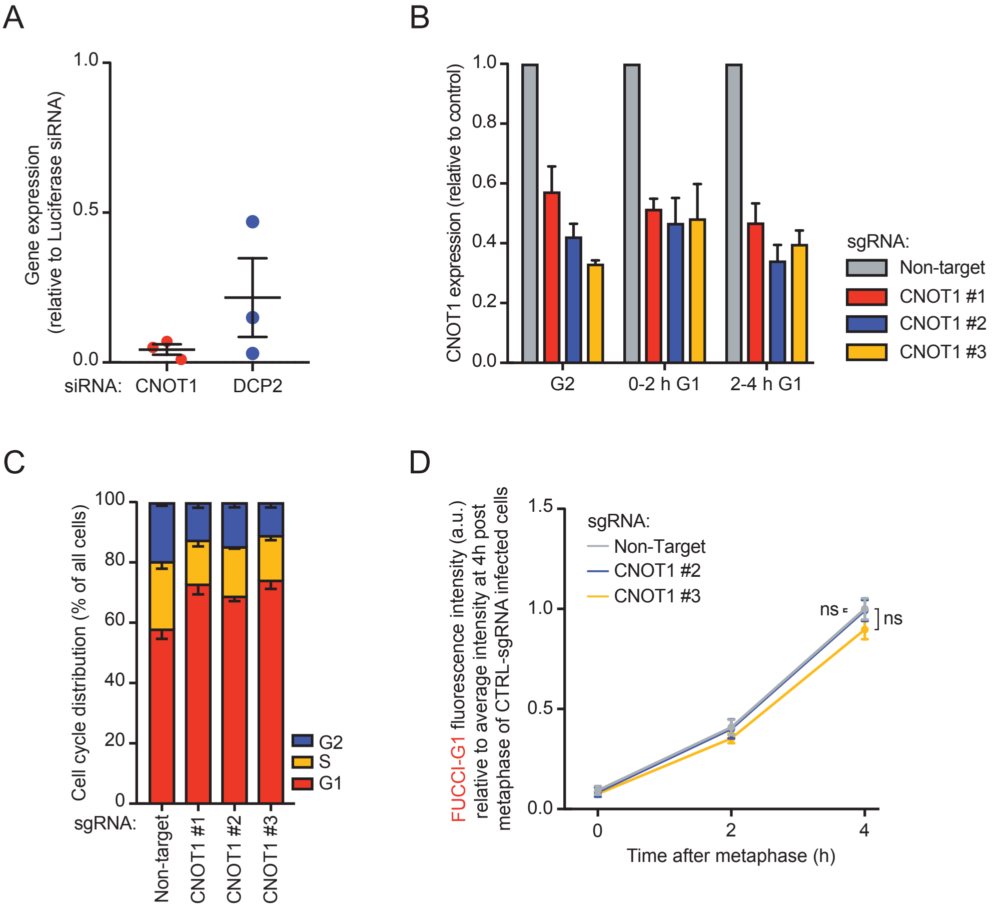
