## Supplemental Figure 2 for "Time-resolved single-cell sequencing identifies multiple waves of mRNA decay during mitotic exit"

### Figure S2 - Supplementary to Figure 2.

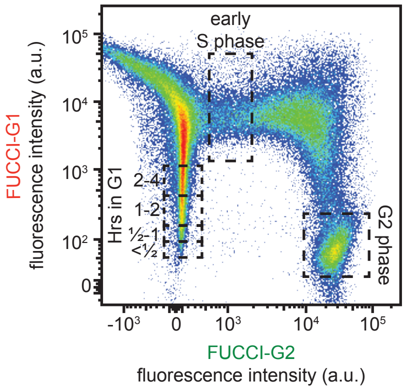


| | |
|---|---|
| Mean fluorescence intensity (a.u.) of S phase gate | 6517 |

| Time in G1 phase | Fluo. int. relative to S phase (lower/upper) | Sort gate FUCCI-G1 fluorescence int. (lower/upper) |
|---|---|---|
| < ½ hour | ≤ 0.0096 | ≤ 63 |
| ½ - 1 hour | 0.0097 - 0.0272 | 63 / 177 |
| 1 - 2 hours | 0.0273 - 0.0795 | 178 / 518 |
| 2 - 4 hours | 0.0796 - 0.2339 | 519 / 1524 |
